# Comparative analysis of syngeneic mouse models of high-grade serous ovarian cancer

**DOI:** 10.1101/2023.03.09.531888

**Authors:** David P Cook, Kristianne JC Galpin, Galaxia M Rodriguez, Noor Shakfa, Juliette Wilson-Sanchez, Madison Pereira, Kathy Matuszewska, Jacob Haagsma, Humaira Murshed, Alison O Cudmore, Elizabeth MacDonald, Alicia Tone, Trevor G. Shepherd, James J Petrik, Madhuri Koti, Barbara C. Vanderhyden

## Abstract

Ovarian cancers often exhibit high rates of recurrence and poor treatment response. Preclinical models that recapitulate the heterogeneity of human disease are critical to develop new therapeutic approaches. While patient-derived models are a powerful tool for testing various therapeutics, their dependence on immune-compromised mice is severely limiting. Syngeneic mouse models, however, allow for the generation of tumours comprising the full repertoire of non-malignant cell types. Here we have performed a comparative analysis of diverse models of high-grade serous ovarian cancer based on transcriptomic profiling of 22 cell line models, and intrabursal and intraperitoneal tumours from 12 models. Among cell lines, we identify distinct features in signalling activity, such as elevated inflammatory signalling in STOSE and OVE16 models, and MAPK/ERK signalling in ID8 and OVE4 models; metabolic features, such as predicted reduction in glycolysis associated with subsets of engineered ID8 subclones; and relevant functional properties, including differences in EMT activation, PD-L1 and MHC class I expression, and predicted chemosensitivity. Finally, we evaluate variability in properties of the tumour microenvironment among models. We anticipate that this work will serve as a valuable resource, providing new insight to help in the selection of models for specific experimental objectives.

## Introduction

Approximately 1% of North Americans born with ovaries will lose their lives to ovarian cancer. Despite decades of research, the standard treatment for many of these individuals remains unchanged. The discovery of synthetic lethality upon exposure of homologous recombination-deficient (HRD) malignant cells to PARP inhibition provided a targeted therapy for approximately 50% of patients with high-grade serous ovarian cancer (HGSOC)--the most common and most lethal form of ovarian cancer ^1–3^. However, many patients with HRD HGSOC fail to respond to this treatment or develop resistance following prolonged treatment ^4^. Developing targeted therapies for the remaining 50% of patients has been challenging due to both the phenotypic and genetic diversity of malignant populations ^5,6^. This has put an increased demand on preclinical models to faithfully recapitulate this complexity.

Preclinical models for HGSOC are fortunately abundant. Human-derived models sample genetic diversity of the disease ^7,8^, but are incapable of being used to model tumours comprising the complete repertoire of non-malignant stromal cells. Successful culture of tumour explants can maintain malignant-stromal interactions *ex vivo*, but these models are not amenable to long-term propagation ^9,10^. Establishing patient-derived xenografts in humanized mouse models with adoptive transfer of autologous leukocytes restores components of stromal interactions ^11^, but it is unclear how accurately these models mirror the progression and therapy response of the native tumour. Thus, human-derived models may be ideal for cell autonomous properties of the malignant population, but are limited in their ability to model cancer progression and the complex interactions within the tumour microenvironment (TME).

Syngeneic mouse models are powerful resources that allow for the generation of tumours in immunocompetent mice. Their ability to model the genetics of human disease is limited, but gene editing strategies can be used to engineer clinically relevant mutations, such as the nearly ubiquitous *TP53* mutation observed in 96% of HGSOC tumours ^1^. However, the complex genomic rearrangements observed in HGSOC have been more challenging to model. Despite this limitation, orthotopic tumours can be generated from these models with histological features consistent with human disease, making them a prime resource for studying both the evolution of a complex TME within the ovary and testing of therapeutics that depend on interactions within it.

Various syngeneic models of HGSOC have been developed, but the strategies to establish them–the cell type of origin, the strategy for malignant transformation, and the engineering of relevant mutations–have varied. Differences in the experimental progression of these models have been reported, including differences in growth rate, TME composition, and sensitivity to treatment ^12–16^. Thus, there is a need to comprehensively compare these models in order to understand their inherent differences. In this study, we have performed transcriptomic profiling of 22 syngeneic models, as well as tumours derived from select models. We evaluate inherent differences associated with cell type of origin, the phenotypic divergence associated with spontaneous transformation from prolonged *in vitro* culture, and the impact of engineering clinically relevant mutations. We explore how these phenotypes give rise to tumours with unique TMEs, such as intrabursal STOSE tumours with low stromal content or the lymphocyte-rich ID8-*Trp53^-/-^* model. Together, this work provides insight into the properties of diverse syngeneic models that can inform the selection of appropriate models for subsequent research and therapeutic testing.

## Results

### Transcriptomic profiling of syngeneic mouse models of high-grade serous ovarian cancer

To develop a resource of transcriptomic data that could provide further insight into models of HGSOC, we performed RNA-seq on a collection of commonly used mouse ovarian cancer cell lines (**Figure 1A**). This collection comprises models from both oviductal (OVE/MOE) and ovarian surface epithelium (OSE), spontaneously transformed (STOSE and ID8) models, secondary lines derived from ascites, and models engineered with clinically relevant mutations in tumour suppressor genes or constitutive activation of oncogenes. For select models, we also sequenced RNA from tumours derived from either intrabursal (IB) or intraperitoneal (IP) injection of the cells, allowing us to evaluate how properties of the models and TME may affect features of the resultant tumours (**Figure 1A**).

**Figure 1.**
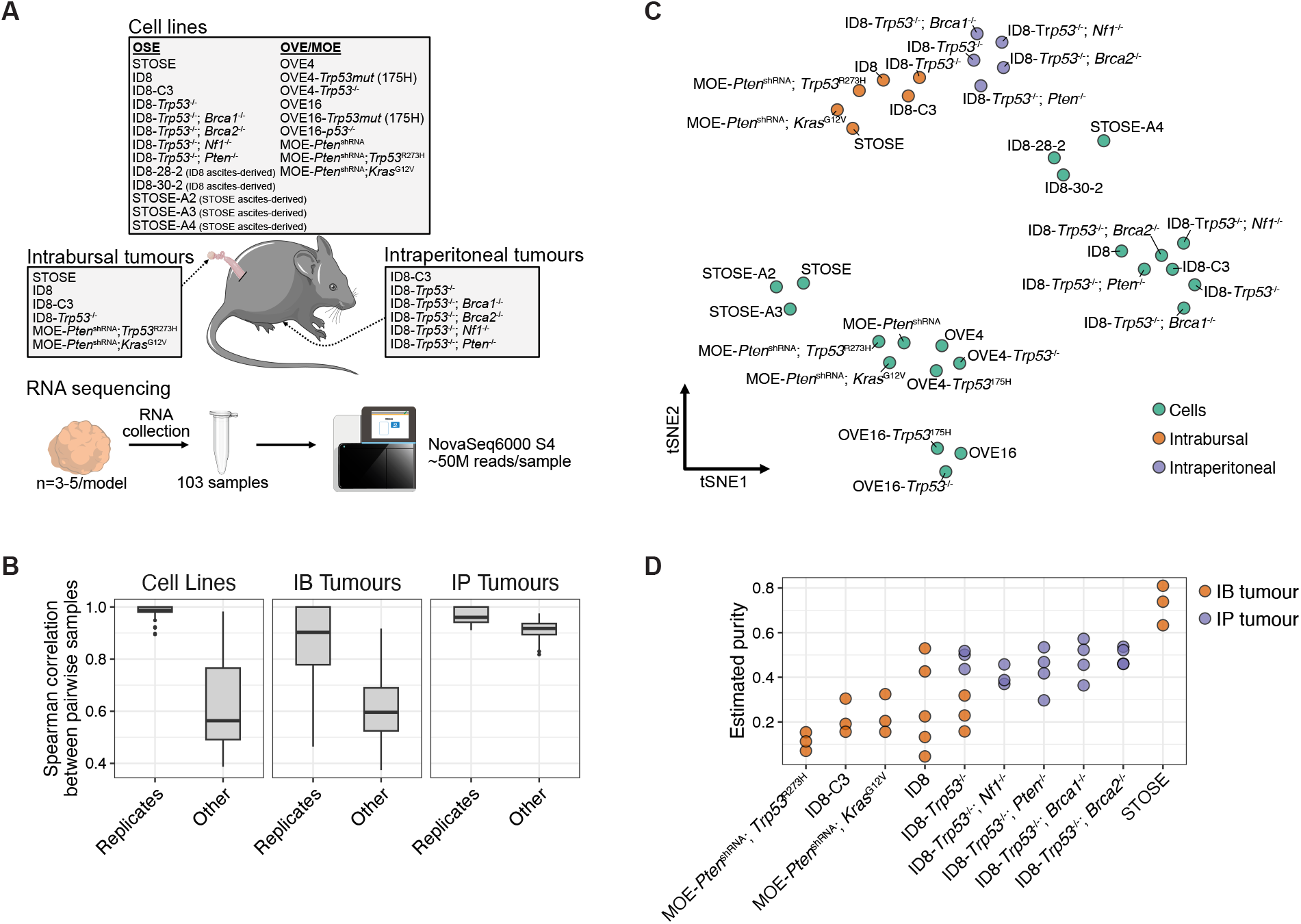
Transcriptional profiling of diverse syngeneic mouse models of HGSOC. **A.** Schematic of the study design. RNA-seq was performed on all listed models and 3-5 replicates were included for each model. **B.** Comparison of spearman correlation coefficients of transcriptional profiles both within and between models for cell lines, IB tumours, and IP tumours. **C**. tSNE embedding of the replicated-averaged transcriptional profile of each model. **D.** Tumour purity estimates from IB and IP tumours based on the estimated proportion of malignant cells predicted by deconvolving bulk RNA-seq profiles with CIBERSORTx. Cell type profiles used from deconvolution were derived from scRNA-seq data of STOSE and ID8 tumours ^13^. **E.** Survival Kaplan-Meier plots of FVB/N (MOE-*Pten*^shRNA^; *Trp53^-/-^*) and C57BL/6 tumor bearing mice (*ID8-Trp53^-/-^*;*Brca2^-/-^*, ID8-*Trp53^-/-^*), treated with 20mg/kg carboplatin (red line) or saline (black dotted line). Curves represents n=10 mice per group per model. Log-rank (Mantel-Cox) \**p* <0.05.

Across all models, we found transcriptional profiles of biological replicates (n=3-5/model) were strongly correlated (**Figure 1B**). Unsurprisingly, bulk profiles from tumour samples shared little similarity to those of pure cell lines (**Figure 1C**). They did, interestingly, cluster according to whether they were generated through IP or IB injection of cells, likely reflecting inherent differences in TMEs of these models. In general, IB tumours were more variable, even among replicates (**Figure 1B**). To evaluate differences in the TME between these two tumour sites, we used CIBERSORTx ^17^ to decompose the bulk RNA-seq profiles and predict the relative proportions of malignant and non-malignant cell types. The inferred contribution of non-malignant stroma to IB tumours was higher than in IP tumours in all models except STOSE IB tumours, which had notably high malignant purity (**Figure 1D**). As such, differences that emerge in each tumour’s microenvironment during its development will have a greater impact on the bulk profiles, contributing to the increased variability among IB samples.

### Cell line models exhibit distinct signalling, metabolic, and functional properties

Principal component analysis (PCA) of the cell line samples highlighted the impact of the models’ cell of origin, with the first principal component (PC) separating OSE- and OVE-derived models (**Figure 2A**). Despite factors such as long-term *in vitro* culture and genetic manipulation, an imprint of the cells’ original identity persisted in culture: various genes uniquely associated with OSE and OVE identity were among the top loaded genes for PC1, including *Pax8* and *Krt7* in OVE lines and *Krt19* and *Amhr2* in OSE-derived lines (**Figure 2B**). We performed gene set enrichment analysis (GSEA) on PC1 loading-ranked genes to identify properties that distinguish these distinct epithelial phenotypes. Consistent with their cell of origin, OVE lines appear more differentiated, with elevated expression of epithelial signatures and MAPK/ERK signalling (**Figure 2C**) ^18^. OSE-derived lines, however, were associated with expression patterns consistent with a more mesenchymal phenotype (**Figure 2C**).

**Figure 2.**
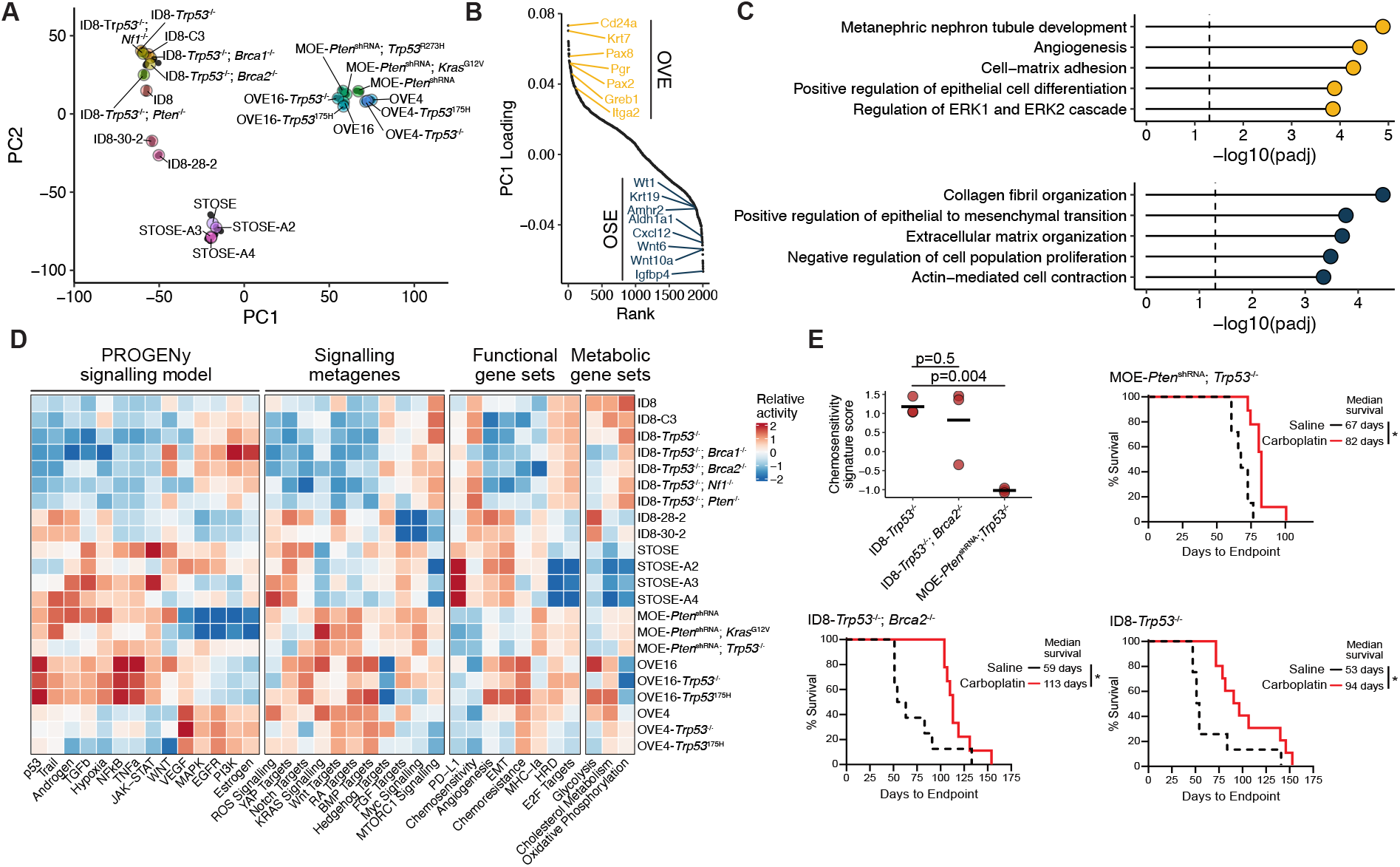
Comparison of cell line models for HGSOC. **A.** PCA embedding of the transcriptional profiles of cell line models. Black points represent individual replicates and coloured points reflect the average coordinates of model replicates. **B.** Ranked gene loadings for the first principal component, which separates OSE and OVE-derived cell lines. Select highly ranked genes are shown. **C.** Over-enrichment analysis of GO terms among the top and bottom 200 ranked genes. P-values were adjusted using the Benjamini-Hochberg method. **D.** Relative (Z-score transformed) gene set scores among cell line models. Relative scores were computed for each replicate prior to averaging scores for each model. The source and genes associated with each gene set are listed in **Supplemental Data 1**.

To further explore biological properties differing among the various models, we collected transcriptional signatures associated with multiple signalling pathways, biological function, and metabolic processes (**Supplemental Data 1**) ^19–24^. We calculated rank-based signature scores for each gene set and compared the relative activity between models (**Figure 2D**). This revealed features that are particularly relevant for the selection of appropriate models for preclinical studies. For example, the ID8 models had higher proliferative activity (E2F targets), with elevated oxidative phosphorylation activity and MTORC1 signalling, but they lacked activity of various cytokine-associated pathways (TGFb, TNFa, NFkB). These features may contribute to a predicted increase in sensitivity to chemotherapy, but these cells also had reduced expression of MHC class I components and expressed higher levels of the immunosuppressive factor PD-L1, which may predict poorer outcomes and/or decreased sensitivity to immunotherapy. In contrast, JAK-STAT, TGFb, and TNF signalling are elevated in OVE16 and STOSE cell lines, which may be ideal to model tumours with high levels of inflammation and chemoresistance (**Figure 2D**). While these inferences are strictly based on expression levels of relevant genes, the predicted decrease in chemosensitivity of the MOE (originally derived from OVE4) cell lines relative to ID8 lines is consistent with survival data from carboplatin-treated IP tumours (**Figure 2E**).

We next examined the impact of p53 mutations on the transcriptional profile of cell lines. We compared the profiles of each cell line harboring either null or oncogenic point mutations to the relevant parental line. These mutations, along with possible divergence following their introduction, resulted in thousands of differentially expressed genes (adj. P-value < 0.05, |log2FC| > 1). Many of these are specific to individual cell lines, though common effects were evident, most commonly between the null and point mutants of the same parental line (**Figure 3A,B).**In general, the OVE-derived models exhibited changes that were more similar to one another than the ID8 line (**Figure 3B**). However, changes associated with the introduction of the *Trp53^-/-^* mutation into the MOE-*Pten*^shRNA^ cell line (originally derived from OVE4 cells) were distinct from those in the OVE4 model (**Figure 3B**). It is unclear if this can be attributed to an interaction between *Pten* suppression and loss of p53, or if it is simply due to divergence of the lines during their manipulation. To assess general features associated with these mutations, we assessed the genes most commonly affected (**Figure 3C**). We found that commonly upregulated genes were often associated with developmental-associated gene sets (eg. axon guidance, integrin-mediated signaling, cartilage development) (**Figure 3D**). Commonly downregulated changes, however, were strongly associated with inflammatory signalling (eg. interferon beta response, chemokine signalling), which may dampen the immune response within the TME (**Figure 3D**).

**Figure 3.**
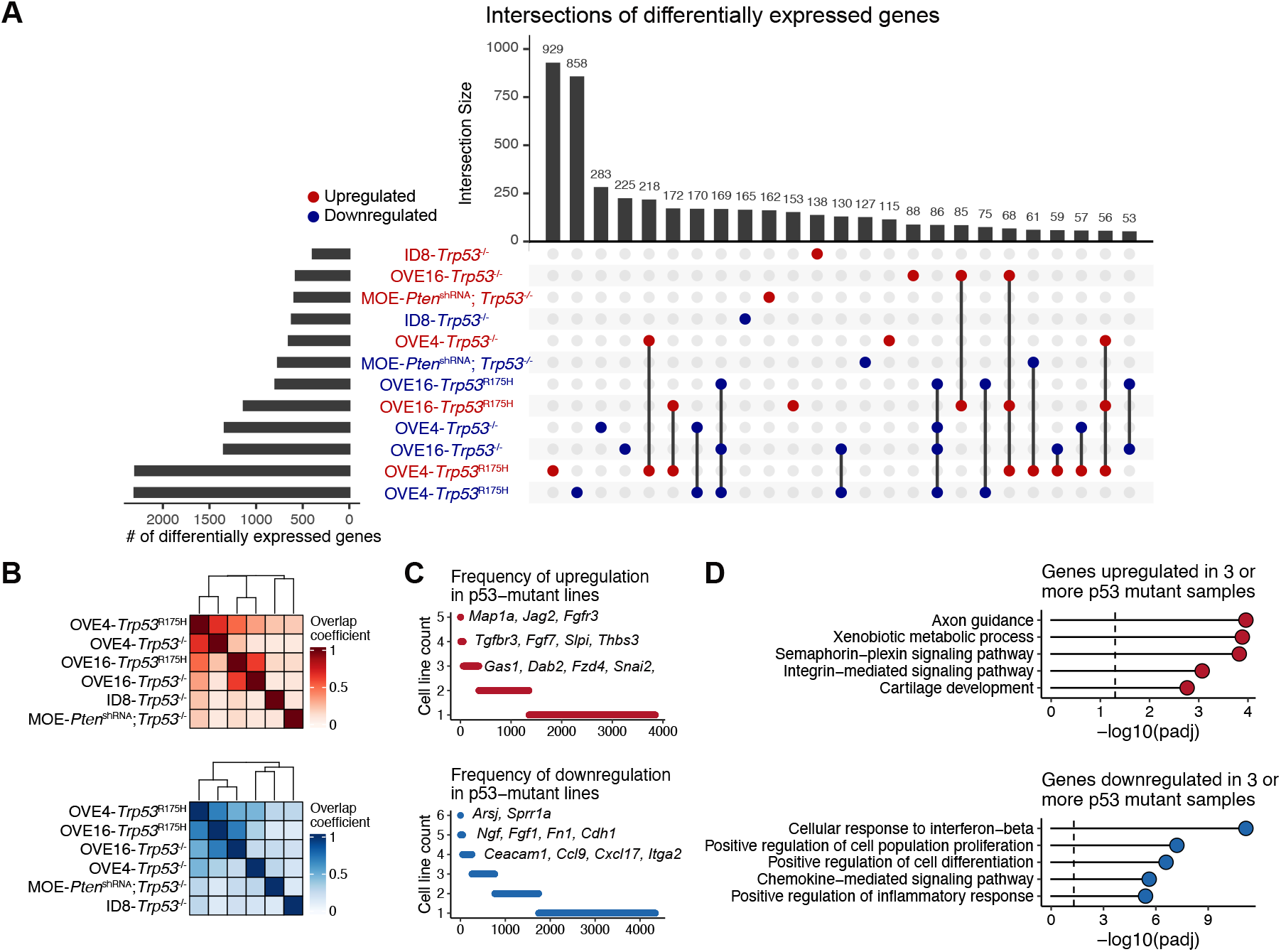
Common and divergent effects of null and point p53 mutations. **A.** Intersections of differentially expressed genes associated with p53 mutations. Red and blue text and point reflect up and down regulated genes, respectively. The top bar chart shows the number of intersecting genes between the conditions highlighted by connected points below. The number of differentially expressed genes per condition is shown on the left. **B.** Clustered heatmap of overlap coefficients associated with up (top) and downregulated (bottom) genes. **C.** The frequency of cell lines in which genes are differentially expressed. Select commonly affected genes are highlighted. **D**. GO terms associated with genes up (top) and downregulated (bottom) in 3 or more cell lines.

### Spontaneous transformation leads to highly divergent phenotypes in STOSE and ID8 models

The STOSE and ID8 models of HGSOC were both derived from prolonged culture of OSE cells that led to progressive aneuploidy and, ultimately, the capacity to form malignant tumours ^25,26^ (**Figure 4A**). It has been demonstrated that these models produce tumours with distinct microenvironments ^13^, but beyond each model having a unique constellation of genomic aberrations, the phenotypic differences of these lines have yet to be characterized. To begin to address this, we evaluated genes differentially expressed in the STOSE and ID8 cell lines. Despite being derived from the same cell type–albeit different mouse strain–we observed massive divergence in their phenotypes, with 5115 differentially expressed genes (p < 0.05; |logFC| > 1) (**Figure 4B**). We evaluated GO terms enriched in the genes associated with each line and found that STOSE were associated with mesenchymal [extracellular matrix (ECM) organization, migration] and immunoregulatory terms, whereas ID8 cells were primarily characterized by higher expression of various metabolic pathway components (**Figure 4C**).

**Figure 4.**
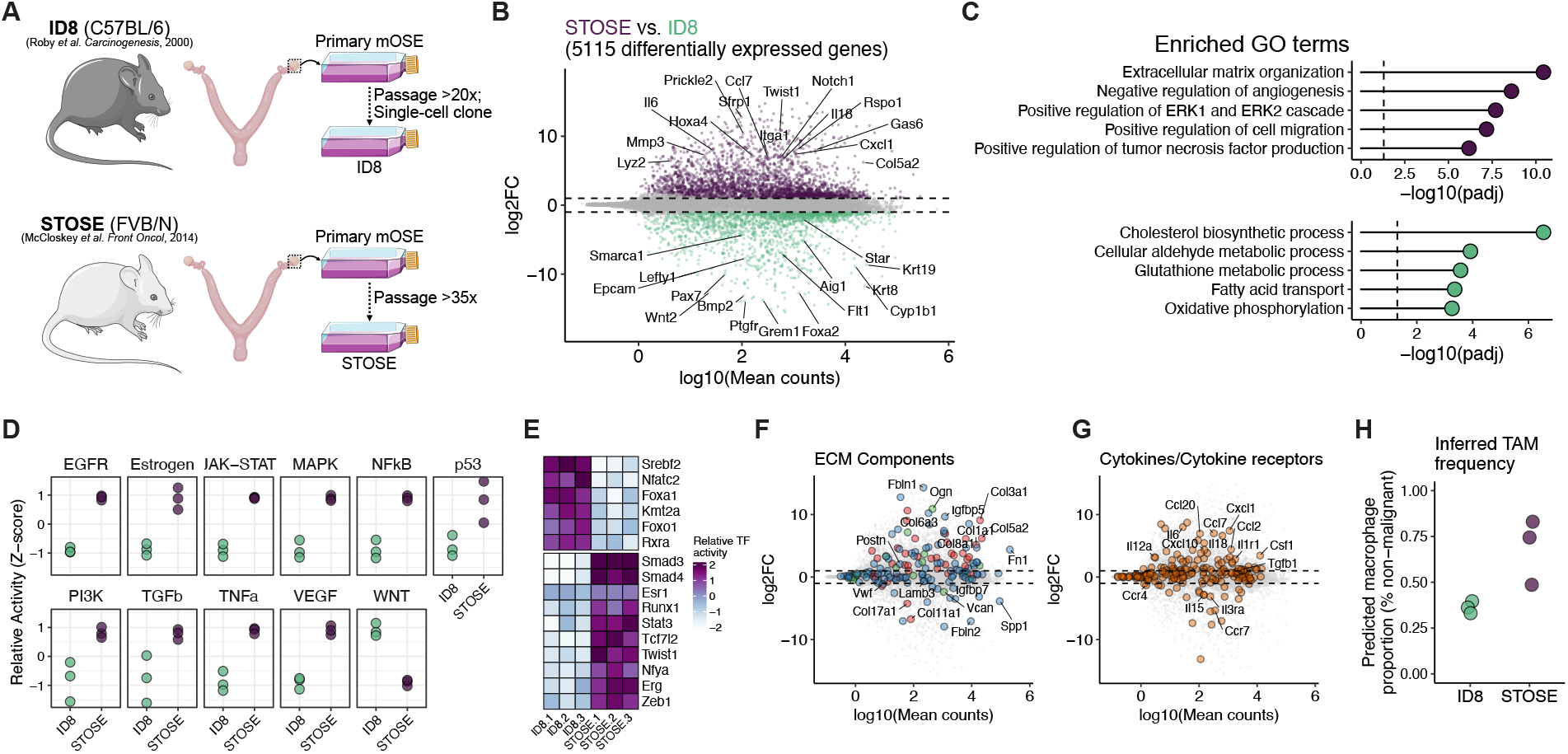
Divergence of spontaneously transformed ovarian surface epithelium. **A.** Schematic demonstrating the establishment of two spontaneously transformed models of HGSOC (STOSE and ID8). **B.** Differentially expressed genes between STOSE and ID8 cells. The x-axis reflects the average quantification (mean counts) for each gene and the y-axis shows the log2 fold change (log2FC) between the two models. The dashed line reflects a |log2FC| threshold of 1 and only genes with an adjusted p-value < 0.05 are shown. Select genes are labelled. **C.** Over-representation analysis of significant genes associated with each cell line. **D.** Inferred activity of pathways predicted to be differentially active between the two models. Activity scores were calculated using PROGENy ^19^. **E.** Predicted Inferred activity of transcription factors predicted to be differentially active between the two models. Activity scores were calculated based on regulons from DoRothEA ^41^. **F.** Identical plot to (B), highlighting collagens (red), glycoproteins (blue), and proteoglycans (green) from the MatrisomeDB^42^. **G.** Identical plot to (B), highlighting cytokines and their receptors from the KEGG pathway “Cytokine-cytokine receptor interaction”. **H.** Predicted proportion of tumour-associated macrophages (TAMs) among the non-malignant fraction of IB tumours. Proportions were based on bulk RNA-seq deconvolution using CIBERSORTx.

Given the challenges of interpreting gene set over-representation with such disparate expression profiles, we evaluated several specific properties of these cells. First, we inferred signalling and transcription factor activity in the two models. Consistent with the mesenchymal and immunoregulatory GO term enrichment, conserved targets of many signalling pathways were preferentially activated in STOSE, including TGFb, TNFa, NFkB, and MAPK signalling (**Figure 4D**). Related transcription factors (eg. Smad3/4, Stat3, Twist1, Zeb1) were also predicted to have higher activity in STOSE (**Figure 4E).** Evidence of signalling patterns specific to ID8 cells was limited. Wnt targets were more highly expressed, along with transcriptional activity of the epithelium-associated transcription factors Foxa1 and Foxo1, and Srebf2, which controls cholesterol homeostasis, consistent with metabolic changes we observed more globally (**Figure 4E**).

The consistent observation of mesenchymal properties in STOSE cells led us to evaluate the expression of ECM components in the two models (**Figure 4F**). Many components were reliably detected but were not differentially expressed between the two models. However, STOSE cells had elevated expression of a variety of collagens (*Col1a1*, *Col5a3*, *Col4a4*, *Col3a1*, and others) and ECM glycoproteins, including the canonical mesenchymal marker *Fn1*. Notably fewer ECM components were preferentially expressed in ID8 cells (eg. *Col11a1*, *Col17a1*, *Igfbp7*, *Fbln2*) (**Figure 4F**)

To further explore differences in the expression of immunoregulatory factors, we evaluated the expression of cytokines, chemokines, and their receptors in the two models (**Figure 4G**). Consistent with GO term enrichment, STOSE cells expressed higher levels of diverse chemokines and cytokines, including *Il6*, *Il18*, *Tgfb1*, *Ccl7*, and *Csf1*. Similar to ECM components, fewer factors were more dominantly expressed in ID8 cells. They did, however, express higher levels of *Il15*, *Cxcl16*, and various cytokine/chemokine receptors (*Ccr7*, *Ccr4*,and *Il3ra*) (**Figure 4G**). We predicted that the imbalanced expression of immunoregulatory factors may lead to differences in the immune infiltration. Using the estimated cell type proportions from the deconvolution of bulk RNA-seq from the tumours of these cells, we found that intrabursal injection of STOSE cells consistently yielded higher purity tumours with a reduced proportion of immune cell infiltration (**Figure 1D**). Among the non-epithelial fraction in both models, macrophages were more prevalent in STOSE (**Figure 4H**), consistent with their elevated chemokine expression. These inferences from bulk RNA-seq deconvolution are consistent with recent flow cytometry-based immunophenotyping and cytokine profiling of these models ^13^.

### Secondary cell lines derived from ascites stably activate mesenchymal programs

Collecting metastatic cells from mouse tumour models for the purpose of deriving aggressive sub-lines is a common strategy. Tumours from these sub-lines often progress more rapidly and metastasize more consistently than parental models. In our cohort of samples, we have included ascites-derived lines from both STOSE and ID8 intrabursal tumours to evaluate how they deviate from their parental lines and to determine if independent lines derived from ascites acquire common traits (**Figure 5A**).

**Figure 5.**
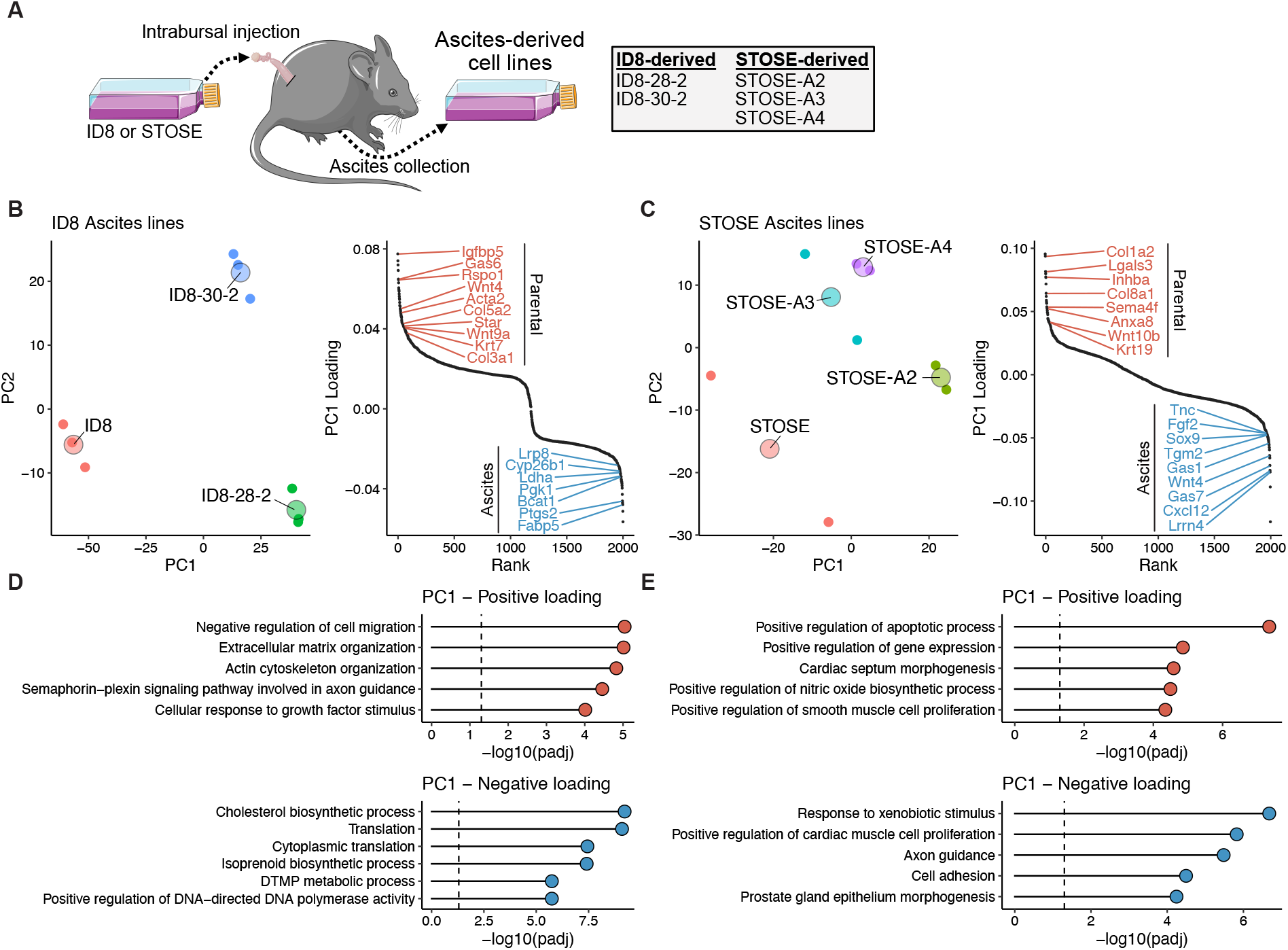
Activation of EMT-associated characteristics in ascites-derived lines. **A.** Schematic of ascites-derived subclones of ID8 and STOSE models. **B**. PCA embedding (left) and ranked gene loadings for PC1 (right) for the ID8 models. **C.** Same as (B) for STOSE-derived models. **D.** Over-representation analysis of GO terms associated with the top and bottom 200 loaded genes for PC1 in the ID8 model. **E.** Same as (D) but for STOSE-derived models.

We used PCA to evaluate the relationship between ascites lines and their parental cell lines. For both STOSE and ID8 lines, the first PC reflected a differentiation axis that separated ascites-derived lines from the parental model (**Figure 5B,C**). Ascites lines from the ID8 model (28-2, 30-2) lose activity in biosynthetic pathways (eg. cholesterol, ribogenesis, and isoprenoid pathways), activating canonical EMT features, including signatures associated with ECM organization, cytoskeleton remodelling, and TGFb signalling (**Figure 5D**). Notably, there was no particularly striking reduction in epithelial features in these ascites lines, consistent with a hybrid epithelial-mesenchymal phenotype ^18,27^. The activation of EMT features in STOSE ascites was less evident, perhaps due to the more-mesenchymal features of the parental lines, but a suppression of genes associated with epithelial morphogenesis was a prominent trend (**Figure 5E**). Despite resulting in more aggressive tumours, markers of proliferation were less abundant in ascites lines from both models (E2F Targets; **Figure 2D**).

### Engineering mutations in tumour suppressor genes produces novel phenotypes that affect the TME

Despite modelling various histological and molecular features of HGSOC, STOSE and ID8 models lack mutations commonly observed in the disease, such as *Trp53*, which is mutated in 96% of HGSOC cases ^1^. To better recapitulate the genetics of the disease, various clinically relevant mutations have been engineered into the ID8 model using the CRISPR-Cas9 system ^28,29^. In our cohort of samples, we have included the ID8-Trp53*-/-* model along with four lines with a second inactivating mutation in either *Brca1* (23% of HGSOC cases), *Brca2* (11%), *Nf1* (12%), or *Pten* (7%) ^1^ (**Figure 6A**).

**Figure 6.**
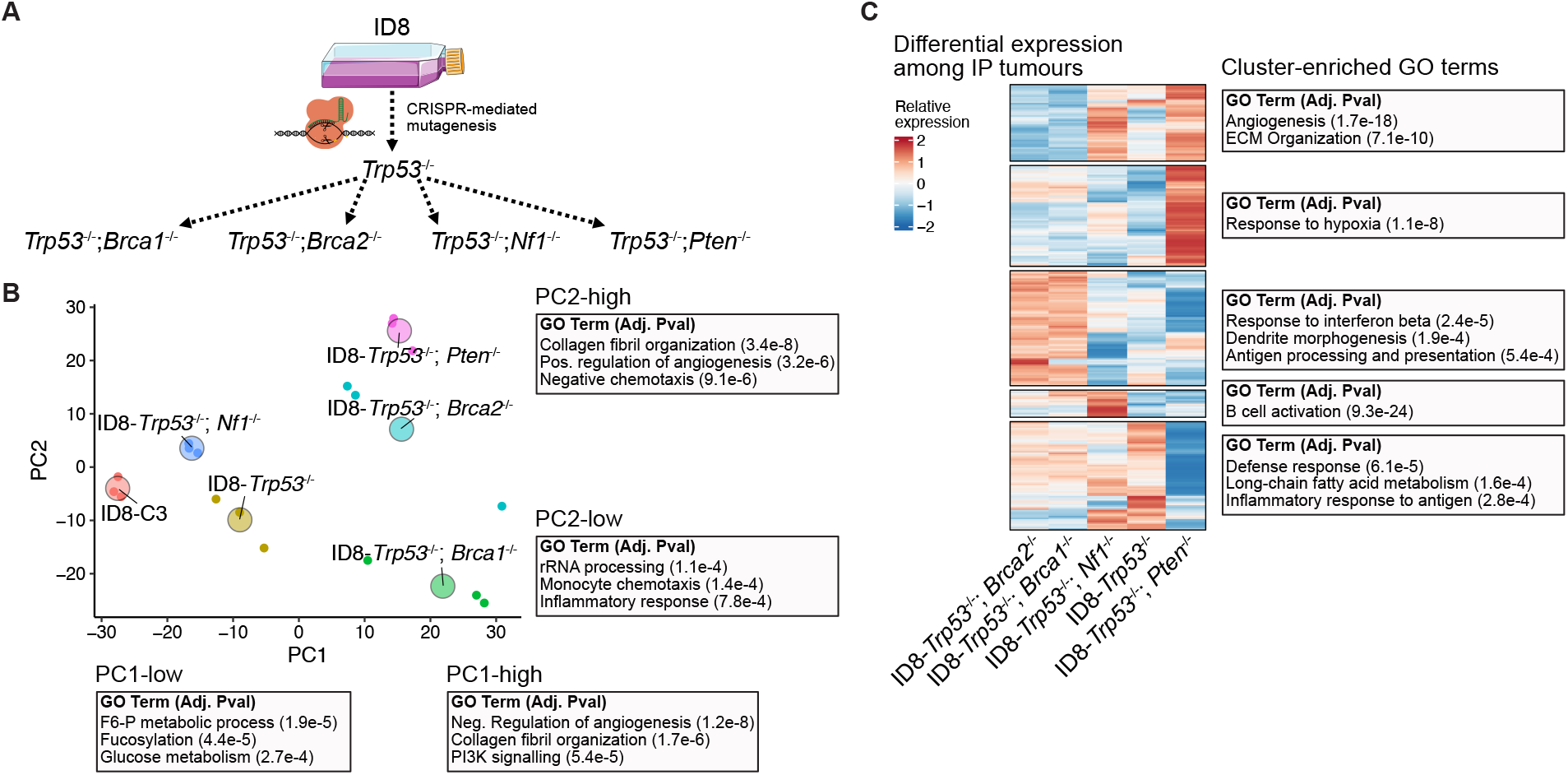
Variation among IP tumours of the ID8 model engineered with clinically relevant mutations. **A.** Schematic of the mutant ID8 lines. All mutations were previously engineered with CRISPR-mediated mutagenesis^28,29^. **B.** PCA embedding of the transcription profile from bulk IP tumour samples. Small points represent individual replicates of each model and the large labelled point is at the average coordinates for each model. Over-represented GO terms among top and bottom loaded genes for both PC1 and PC2 are shown next to the embedding along with Benjamini-Hochberg-adjusted p-values. **C.** Clustered heatmap of the 1544 genes differentially expressed (likelihood ratio test; adj. p-value < 0.05) among tumours from the ID8 models. Selected enriched GO terms for each cluster are shown next to each cluster.

PCA of the cell line gene expression profiles highlighted the phenotypic divergence following introduction of tumour suppressor mutations (**Figure 6B**). Similar to the analysis of ascites-derived lines, the first PC seemed to reflect an axis of differentiation, with control, *Trp53-/-*, and *Trp53-/-*;*Nf1-/-* ID8 cells having evidence of elevated cholesterol biosynthesis and fructose 6-phosphate metabolism. The expression of these factors is suppressed in the *Trp53-/-*;*Pten-/-*, *Trp53-/-*;*Brca1-/-*, and *Trp53-/-*;*Brca2-/-* cells, which preferentially activate mesenchymal signatures (**Figure 6B**). Among these less differentiated lines, we found that the *Trp53-/-*;*Brca1-/-* model expressed higher levels of monocyte chemotaxis factors whereas the *Trp53-/-*;*Pten-/-* model expressed inhibitors of chemotaxis, along with various ECM remodelling factors (collagen organization, positive regulation of angiogenesis) (**Figure 6B**).

We next evaluated whether these features may lead to distinct features in the TME of each model. Evaluating the bulk RNA-seq profile from whole tumour lysate for each model, we identified genes differentially expressed among the models and clustered genes based on their expression across models. Many clusters contained markers highly specific for non-malignant cell types, reflecting differences in the relative abundance of those cell types across models (**Figure 6C**). Generic immune/defense signatures–likely associated with total immune infiltration–were notably deficient in the *Pten^-/-^* models (**Figure 6C**). The *Brca1^-/-^* cell line expressed various monocyte recruitment factors *in vitro* and the *Brca1^-/-^* and *Brca2^-/-^* tumours had evidence of antigen processing expression in their bulk RNA profiles (**Figure 5C**). Increased chemokine expression and macrophage infiltration in BRCA-mutant tumours is consistent with recent observations following the addition of a *Brca1^-/-^* mutation to an OVE-derived model engineered with *Trp53*^R172H^, *Pten^-/-^*, *Nf1^-/-^*, and *Myc^OE^* mutations ^15^. *Pten^-/-^* tumours, and to a lesser extent the *Nf1^-/-^* model, had increased expression of angiogenic signatures, further consistent *in vitro* expression patterns (**Figure 6C**). Finally, the *Nf1^-/-^* model had a unique enrichment of diverse immunoglobulins, reflecting an abundance of B cells (**Figure 6C**).

## Discussion

Preclinical models are an invaluable resource to further our understanding of ovarian cancer, enabling the development of strategies to improve patient care. The number of ovarian cancer models is expanding and it has become increasingly unclear which models are most appropriate for specific experimental objectives. Here, we have focused on cataloging and comparing transcriptomic data from a cohort of syngeneic mouse models of HGSOC.

Just as the molecular features of human tumours are diverse, these models displayed remarkable variation. While there has been a great focus on deriving models with mutations that recapitulate the genetics of human HGSOC, the lack of consistent mutations (beyond *Tp53*) and the diverse copy number alterations observed in human tumours make this goal intractable and perhaps irrelevant. This also raises the question of whether it’s feasible to develop therapies that are selective to the genetics of the ~50% of tumours that are HR proficient. Effort in this direction is certainly critical and it is possible that a collection of targeted therapies could be utilized in conjunction with genetic screening to deliver personalized treatments to these patients. However, the challenges introduced by these complex genetics could be curtailed by focusing on developing therapeutics that are not dependent on them. For example, perturbations to signalling pathways may be used to sensitize chemoresistant phenotypes ^30^, or immunotherapies may be delivered to rejuvenate immune activity within the TME. Certainly there has been increased effort in this direction over the recent years. Therefore, there is an urgency to prioritize models based on their ability to recapitulate the general phenotypes and structural properties of human disease rather than the specific genetics.

OSE-derived models have received criticism for not reflecting the likely cell of origin for the majority of HGSOC. While this criticism is valid, it should not discount the information that these models have provided or the applicability of these models in future research. Both STOSE and ID8 are capable of producing ovarian tumours with histological features consistent with human HGSOC ^25,26^, and so they – like all models – exhibit various features that are consistent with human disease, and others that are not. Importantly, they produce tumours with distinct properties (eg. the characteristically low stromal content of STOSE IB tumours), providing an opportunity to test therapeutics against diverse TMEs similar to those seen in human tumours. Similarly, although clinically relevant mutations have been engineered into the ID8 model, it is unclear if the phenotypic differences are strictly related to specific mutations that have been engineered or if they reflect divergence that occurred throughout the engineering process. It is perhaps more relevant that this has produced distinct sub-models with unique TME properties, such as the vascular rich *Pten^-/-^* model or the macrophage-rich *Brca1/2^-/-^*models.

Given the importance of recapitulating the TME of human disease, the experimental strategy for generating tumours is particularly relevant. We demonstrated that orthotopic tumours generated from intrabursal injection have a consistently more abundant tumour stroma than solid masses collected following intraperitoneal injection. This may simply owe to differences in tissue properties (eg. stiffness, adiposity, etc.) between the ovary and sites within the peritoneal cavity (eg. omentum). IP injection is experimentally convenient and often justified as a model of metastatic disease, but it is unclear if the resultant lesions faithfully recapitulate those emerging from the natural metastatic cascade from the ovary to these sites. Specifically, IP injection fails to impose the selective bottleneck that enriches for cells capable of leaving the adnexa. Rather, it gives the entire malignant population the opportunity to seed metastases, including cells that may not be capable of dissemination from a primary tumour. In our analysis, we demonstrated that ascites-derived cells are quite distinct from the parental population. It is unclear if this suggests that phenotypic selection is critical to metastatic dissemination or if the peritoneal cavity simply promotes phenotypic reprogramming.

While we present a relatively large cohort of syngeneic models, this list is by no means exhaustive and further work is required to examine additional models. The addition of other data modalities (eg. epigenetic profiling, deeper genetic characterization, scRNA-seq of tumour models, etc) would also provide valuable information about these models. Together, these initial comparisons have provided insight into phenotypic differences between the various models that ultimately affect the properties and progression of the tumours they make. Further validation of specific functional properties is certainly critical; however, we anticipate that this data will be particularly useful to aid in the selection of appropriate models for specific experimental aims.

## Methods

### Cell lines

Unmodified ID8 cells were provided by Kathy Roby (Roby et al., 2000). ID8-*Trp53^-/-^* F3, ID8-*Trp53^-/-^Brca1^-/-^*, ID8-*Trp53^-/-^Brca2^-/-^*, ID8-*Trp53^-/-^Nf1^-/-^*, ID8-*Trp53^-/-^Pten^-/-^*, were generated by CRISPR-Cas9 mediated knockout, and ID8-C3 (CRISPR control) were generously provided by Dr. Iain McNeish (Imperial College London) ^28,29^. ID8-WT and its derivatives were maintained in Dulbecco’s Modified Eagle Medium (DMEM, Corning) + 4% fetal bovine serum (FBS, Hyclone) and 0.01 mg/mL insulin-transferrin-sodium-selenite solution (ITSS; Roche) as previously described ^28,29^. ID8 ascites-derived lines (28-2, 30-2) were generated by culturing adherent cells from ascites formed in tumour-bearing mice approximately 60 days following orthotopic injection of parental ID8 cells.

STOSE cells were generated previously ^26^ and STOSE ascites-derived (STOSE-A2, A3, A4) cells were derived from ascites collected from three STOSE-tumour bearing mice. STOSE and STOSE-A cell lines were maintained in a-MEM (minimal essential media) + 4% FBS and ITSS and 2μg/ml epithelial growth factor (EGF, R&D). OVE4 and OVE16 were oviductal epithelial cell lines generated previously ^31^. OVE4 were modified by Dr. Joanna Burdette (Univeersity of Illinois, Chicago) to generate *MOE-Pten^shRNA^*, *MOE-Pten^shRNA^*:*Trp53^R273H^*, and MOE-*Pten^shRNA^Kras^G12V^*, and were maintained in a-MEM + 4% FBS, ITSS, 2μg/ml EGF, and 18.2 ng/mL β-estradiol as previously described ^32,33^. OVE4 and OVE16 were transduced with a lentival vector to expression *Trp53*^R175H^ to generate OVE4-*Trp53*^175H^ and OVE16-*Trp53*^175H^ cell lines. To generate *OVE4-Trp53^-/-^* and OVE16-*Trp53^-/-^* lines, OVE cells were transfected with pSpCas9-Trp53 plasmids using Lipofectamine and subjected to puromycin selection. *Trp53* deletion and the R175H mutation were validated by Western blot. Mycoplasma testing was performed prior to animal experiments. Cells were incubated at 37°C with 5% carbon dioxide

### Generation of tumours with syngeneic models

Animal experiments to generated orthotopic tumours were carried out using protocols approved by the Animal Care Committee at the University of Ottawa and conforming to the standards defined by the Canadian Council on Animal Care (CCAC). FVB/N mice (for STOSE and MOE cell lines) were acquired from Charles River Laboratories and C57BL/6 mice (for ID8 and derivatives) were purchased from The Jackson Laboratory. Orthotopic intrabursal (IB) tumours were generated by injecting 1.5 ×10^5^ cells (ID8-WT, ID8-C3,ID8-*p53^-/-^*, STOSE, MOE-PTEN/p53, or MOE-PTEN/KRAS) under the ovarian bursa in 2μl phosphate-buffered saline (PBS) as previously described ^26^. Primary tumours were collected when mice reached humane endpoint, snap-frozen in liquid nitrogen, and stored at −80 degrees Celsius until RNA extraction.

Procedures to generate IP tumours were approved by the University Animal Care Committee (UACC) at Queen’s University and the CCAC. The various modified ID8 cell lines (5-6 × 10^6^, n=5-10 per genotype) in 350 μL of PBS were transplanted via intraperitoneal injections into C57BL/6 female mice aged 8 to 10 weeks (Charles River Laboratories).

All mice were maintained in specific pathogen-free conditions. Mice were left untreated until they reached their endpoint, which for IP tumours was deemed the point at which their abdominal diameters were approximately 34 mm. The tumours were snap-frozen in liquid nitrogen and stored at −80°C until RNA extraction.

### In vivo carboplatin sensitivity

Mice were injected intraperitoneally with 5×10^6^ tumor cells resuspended in 100μl PBS. Then, after 25% of tumor establishment for each murine model, mice received bi-weekly injections of carboplatin at 20mg/kg per mouse for a total of 4 weeks (8 doses). Control groups received saline. Mice were followed-up for survival assessment until they reach humane endpoint.

### RNA collection and library preparation

Total RNA was extracted according to the manufacturer’s instructions with the RNeasy Plus Mini Kit (Qiagen) for the majority of samples. IP tumours collected from mice injected with *ID8-Trp53*-/-, *ID8-Trp53*-/-; *Brca1*-/-, *ID8-Trp53*-/-; *Brca2*-/-, *ID8-Trp53*-/-; *Nf1*-/-, and ID8-*Trp53*-/-; *Pten*-/- cells were isolated at endpoint using the total RNA Purification Kit (Norgen Biotek Corporation, ON, Canada) as per the manufacturer’s instructions.

RNA-Seq libraries were generated from 250 ng of total RNA as following: mRNA enrichment was performed using the NEBNext Poly(A) Magnetic Isolation Module (New England BioLabs). cDNA synthesis was achieved with the NEBNext RNA First Strand Synthesis and NEBNext Ultra Directional RNA Second Strand Synthesis Modules (New England BioLabs). The remaining steps of library preparation were done using and the NEBNext Ultra II DNA Library Prep Kit for Illumina (New England BioLabs). Adapters and PCR primers were purchased from New England BioLabs.

The libraries were normalized and pooled and then denatured in 0.02N NaOH and neutralized using HT1 buffer. The pool was loaded at 200pM on a Illumina NovaSeq S4 lane using Xp protocol as per the manufacturer’s recommendations. The run was performed for 2×100 cycles (paired-end mode). A phiX library was used as a control and mixed with libraries at 1% level. Base calling was performed with RTA v3. Program bcl2fastq2 v2.20 was then used to demultiplex samples and generate fastq reads

### RNA-seq transcript quantification and processing

Pseudoalignment and transcript quantification for each sample was performed using Kallisto (v0.45.0) ^34^ with the GRCm38 build of the mouse genome. The R package tximport (v1.24.0) was used to load transcript quantifications, converting to gene-level transcript estimates. Principal component analysis (PCA) was performed on zero-centered normalized counts from DESeq2’s variance stabilizing vst() transformation. For each independent PCA, only the top 2000 variable genes were used as input. A t-distributed stochastic neighbor embedding (tSNE) was generated for the whole dataset using the top 10 principal components as input into the Rtsne() function provided by the Rtsne R package (v0.16).

### Differential gene expression

All differential gene expression analysis was performed using DESeq2 (v1.36.0) ^35^. To compare STOSE and ID8 cell lines, the Wald test was used to compute p-values and log fold change shrinkage was performed using the apeglm estimator ^36^. To evaluate differentially expressed genes among more than two samples (eg. the collection of modified ID8 lines), a likelihood ratio test was used. Genes with a p-value less than 0.05 and a standard deviation across all tested samples of greater than 0.5 (effectively providing an effect size threshold) were then clustered based on their relative expression levels across samples.

### GO Term over-representation analysis

Over-representation analysis of GO terms among differentially expressed genes was performed using the topGO (v.2.48.0) wrapper topGOtable() provided in the pcaExplorer R package (v2.22.0) ^37^. The elim method implemented in topGO was used to reduce redundancy in the list of enriched gene sets.

### Gene set scoring and inference of signalling and transcription factor activity

The R package singscore (v1.16.0) ^38^ was used to compute gene set activity scores for individual samples. Scores reflect a rank-based statistic from genes comprising each set similar to the Wilcoxon rank sum test. Relative scores between samples were calculated by standardizing scores across samples with a Z-score transformation.

Signalling activity for STOSE and ID8 models was calculated using the R package PROGENy (v1.18.0) ^19^ based on the package’s pretrained regression models of gene activity associated with 14 different signalling pathways. The top 500 genes of each model were used for calculating scores.

Transcription factor activity was inferred using the database of transcription factor targets in the DoRothEA package (v1.8.0) ^39^, only including associations with a confidence level of “A” or “B”. The viper method (v.1.30.0) ^40^ was used to compute activity scores for each individual sample. To compare activities between STOSE and ID8 samples, a simple linear model was used for each factor. Only factors with a Benjamini-Hochberg-adjusted p-value of less than 0.05 were used.

### Bulk RNA-seq deconvolution using scRNA-seq

Cell type deconvolution was performed using a publicly available reference scRNA-seq data of primary orthotopic tumours from both STOSE and ID8 models ^13^ (NCBI GEO Accession: GSE183368). The annotated scRNA-seq data was first used to generate cell type references with CIBERSORTx ^17^, which was then used to deconvolve the normalized (TPM) quantifications for each bulk RNA-seq tumour sample using the default parameters. Predicted cell type proportions were then compared between samples.

## Acknowledgements

Funding was provided by Health Canada to Ovarian Cancer Canada in support of the OvCAN research initiative. We would like to thank Genome Quebec for their assistance in generating the RNA-seq libraries.

## Author Contributions

DPC, AT, JP, MK, and BCV conceived the study. KG, GMR, NS, JWS, MP, KM, JH, and TGS, performed cell culture experiments and collected RNA for library preparation. KG, GMR, EM, HM, NS, JWS, MP, AOC, and KM performed animal studies. DPC performed all computational analysis and drafted the manuscript. All authors revised and finalized the manuscript.

## Data Availability

All RNA-seq data is available at the NCBI GEO Accession GSEXXXXXX.

## Notes

### Competing Interest Statement

The authors have declared no competing interest.

### Summary of Updates

Spelling error in the author list has been corrected. No changes to the manuscript.

## References

1. The Cancer Genome Atlas Research Network. Integrated genomic analyses of ovarian carcinoma. Nature 474, 609–615 (2011).

2. Farmer, H. et al. Targeting the DNA repair defect in BRCA mutant cells as a therapeutic strategy. Nature 434, 917–921 (2005).

3. Bryant, H. E. et al. Specific killing of BRCA2-deficient tumours with inhibitors of poly(ADP-ribose) polymerase. Nature 434, 913–917 (2005).

4. Miller, R. E., El-Shakankery, K. H. & Lee, J.-Y. PARP inhibitors in ovarian cancer: overcoming resistance with combination strategies. J. Gynecol. Oncol. 33, e44 (2022).

5. Cook, D. P. & Vanderhyden, B. C. Ovarian cancer and the evolution of subtype classifications using transcriptional profiling†. Biol. Reprod. 101, 645–658 (2019).

6. Macintyre, G. et al. Copy number signatures and mutational processes in ovarian carcinoma. Nat. Genet. 50, 1262–1270 (2018).

7. Kopper, O. et al. An organoid platform for ovarian cancer captures intra- and interpatient heterogeneity. Nat. Med. 25, 838–849 (2019).

8. Hill, S. J. et al. Prediction of DNA Repair Inhibitor Response in Short-Term Patient-Derived Ovarian Cancer Organoids. Cancer Discov. 8, 1404–1421 (2018).

9. Abreu, S. et al. Patient-derived ovarian cancer explants: preserved viability and histopathological features in long-term agitation-based cultures. Sci. Rep. 10, 19462 (2020).

10. Neal, J. T. et al. Organoid Modeling of the Tumor Immune Microenvironment. Cell 175, 1972–1988.e16 (2018).

11. Odunsi, A. et al. Fidelity of human ovarian cancer patient-derived xenografts in a partially humanized mouse model for preclinical testing of immunotherapies. Journal for ImmunoTherapy of Cancer vol. 8 e001237 Preprint at https://doi.org/10.1136/jitc-2020-001237 (2020).

12. McCloskey, C. W., Rodriguez, G. M., Galpin, K. J. C. & Vanderhyden, B. C. Ovarian Cancer Immunotherapy: Preclinical Models and Emerging Therapeutics. Cancers 10, (2018).

13. Rodriguez, G. M. et al. The Tumor Immune Profile of Murine Ovarian Cancer Models: An Essential Tool for Ovarian Cancer Immunotherapy Research. Cancer Research Communications 2, 417–433 (2022).

14. Shakfa, N. et al. PTEN and BRCA1 tumor suppressor loss associated tumor immune microenvironment exhibits differential response to therapeutic STING pathway activation in a murine model of ovarian cancer. bioRxiv 2022.06.27.497846 (2022) doi: 10.1101/2022.06.27.497846.

15. Iyer, S. et al. Genetically Defined Syngeneic Mouse Models of Ovarian Cancer as Tools for the Discovery of Combination Immunotherapy. Cancer Discov. 11, 384–407 (2021).

16. Zhang, S. et al. Genetically Defined, Syngeneic Organoid Platform for Developing Combination Therapies for Ovarian Cancer. Cancer Discov. 11, 362–383 (2021).

17. Newman, A. M. et al. Determining cell type abundance and expression from bulk tissues with digital cytometry. Nat. Biotechnol. 37, 773–782 (2019).

18. Auersperg, N., Maines-Bandiera, S. L., Dyck, H. G. & Kruk, P. A. Characterization of cultured human ovarian surface epithelial cells: phenotypic plasticity and premalignant changes. Lab. Invest. 71, 510–518 (1994).

19. Schubert, M. et al. Perturbation-response genes reveal signaling footprints in cancer gene expression. Nat. Commun. 9, 1–11 (2018).

20. Liberzon, A. et al. The Molecular Signatures Database (MSigDB) hallmark gene set collection. Cell Syst 1, 417–425 (2015).

21. Wang, Y. et al. Comprehensive Molecular Characterization of the Hippo Signaling Pathway in Cancer. Cell Rep. 25, 1304–1317.e5 (2018).

22. Han, L. et al. Single cell transcriptomics identifies a signaling network coordinating endoderm and mesoderm diversification during foregut organogenesis. Nat. Commun. 11, 1–16 (2020).

23. Sun, J. et al. Large-scale integrated analysis of ovarian cancer tumors and cell lines identifies an individualized gene expression signature for predicting response to platinum-based chemotherapy. Cell Death Dis. 10, 1–12 (2019).

24. Peng, G. et al. Genome-wide transcriptome profiling of homologous recombination DNA repair. Nat. Commun. 5, 1–11 (2014).

25. Roby, K. F. et al. Development of a syngeneic mouse model for events related to ovarian cancer. Carcinogenesis 21, 585–591 (2000).

26. McCloskey, C. W. et al. A new spontaneously transformed syngeneic model of high-grade serous ovarian cancer with a tumor-initiating cell population. Front. Oncol. 4, 53 (2014).

27. Pastushenko, I. & Blanpain, C. EMT Transition States during Tumor Progression and Metastasis. Trends Cell Biol. 29, 212–226 (2019).

28. Walton, J. et al. CRISPR/Cas9-Mediated Trp53 and Brca2 Knockout to Generate Improved Murine Models of Ovarian High-Grade Serous Carcinoma. Cancer Res. 76, 6118–6129 (2016).

29. Walton, J. B. et al. CRISPR/Cas9-derived models of ovarian high grade serous carcinoma targeting Brca1, Pten and Nf1, and correlation with platinum sensitivity. Sci. Rep. 7, 16827 (2017).

30. Boumahdi, S. & de Sauvage, F. J. The great escape: tumour cell plasticity in resistance to targeted therapy. Nat. Rev. Drug Discov. 19, 39–56 (2020).

31. Alwosaibai, K. et al. PAX2 maintains the differentiation of mouse oviductal epithelium and inhibits the transition to a stem cell-like state. Oncotarget 8, 76881–76897 (2017).

32. Eddie, S. L. et al. Tumorigenesis and peritoneal colonization from fallopian tube epithelium. Oncotarget 6, 20500–20512 (2015).

33. Endsley, M. P. et al. Spontaneous Transformation of Murine Oviductal Epithelial Cells: A Model System to Investigate the Onset of Fallopian-Derived Tumors. Front. Oncol. 5, 154 (2015).

34. Bray, N. L., Pimentel, H., Melsted, P. & Pachter, L. Near-optimal probabilistic RNA-seq quantification. Nat. Biotechnol. 34, 525–527 (2016).

35. Love, M. I., Huber, W. & Anders, S. Moderated estimation of fold change and dispersion for RNA-seq data with DESeq2. Genome Biol. 15, 550 (2014).

36. Zhu, A., Ibrahim, J. G. & Love, M. I. Heavy-tailed prior distributions for sequence count data: removing the noise and preserving large differences. Bioinformatics 35, 2084–2092 (2019).

37. Marini, F. & Binder, H. pcaExplorer: an R/Bioconductor package for interacting with RNA-seq principal components. BMC Bioinformatics 20, 331 (2019).

38. Foroutan, M. et al. Single sample scoring of molecular phenotypes. BMC Bioinformatics 19, 404 (2018).

39. Holland, C. H., Szalai, B. & Saez-Rodriguez, J. Transfer of regulatory knowledge from human to mouse for functional genomics analysis. Biochim. Biophys. Acta Gene Regul. Mech. 1863, 194431 (2020).

40. Alvarez, M. J. et al. Functional characterization of somatic mutations in cancer using network-based inference of protein activity. Nat. Genet. 48, 838–847 (2016).

41. Holland, C. H. et al. Robustness and applicability of transcription factor and pathway analysis tools on single-cell RNA-seq data. Genome Biol. 21, 1–19 (2020).

42. Shao, X., Taha, I. N., Clauser, K. R., Gao, Y. (tom) & Naba, A. MatrisomeDB: the ECM-protein knowledge database. Nucleic Acids Res. 48, D1136–D1144 (2019).

